# Improving radio transmitter attachment methods for small mammals through captive trials and field studies

**DOI:** 10.1101/2023.03.08.531796

**Authors:** Freya Robinson, Nikki Van de Weyer, Steve Henry, Lyn A. Hinds, Peter R. Brown, Wendy A. Ruscoe

## Abstract

Radio tracking can be used to collect information about animal movement, home range, behaviour and habitat use. Many field studies have fitted radio transmitters to small rodents using permanent nylon cable tie collars and successfully collected movement and fate data. The approach to animal welfare within the context of scientific research prioritises minimising adverse effects on the research animals. While a range of electronically activated release mechanisms exist in radio tracking collars for larger mammals, weight and size restrictions make these unsuitable for smaller animals (< 30 g). Our aim was to identify a radio transmitter model and attachment method of an appropriate size and weight, which would remain attached to a house mouse (*Mus musculus*) for >20 days to collect movement data and then detach or show signs of detaching after 30 days. Laboratory and field trials established that cable ties with a cotton thread weak-link, using heat shrink to attach a customised radio transmitter worked for wild house mice in agricultural fields. Glue-on methods did not stay attached for long enough to obtain more than a few days tracking data.

**Short summary:** Collecting meaningful radio tracking data for small mammals weighing <30 grams relies on selecting radio transmitter attachment methods suitable for the target species while prioritising animal welfare. Developing a non-permanent radio transmitter attachment for house mice is challenging due to size and weight constraints however, by trialling methods in the laboratory and field we developed a suitable radio collar with an in-built weak-link. Our non-permanent weak-link radio collar is an important improvement on existing permanent radio collars for small mammals.

## Introduction

Radio tracking can be used to collect information about animal movement, home range, behaviour and habitat use, and has been used as a technique in wild animal studies including the understanding of pest house mouse (*Mus musculus*) movement, social structure, habitat use and fate in Australian grain-growing regions (Brown and Singleton 1998; Chambers *et al*. 1999; Krebs *et al*. 1995). Mouse plagues can severely damage crops, livestock and infrastructure, leading to economic loss, health risks and lingering social impacts in crop growing regions (Brown *et al*. 2007; Caughley *et al*. 1994; McLeod 2004). Better understanding of house mouse movement, behaviour and ecology is required to improve management in dynamic cropping systems.

Field studies have fitted radio transmitters to house mice and other small rodents using permanent nylon cable tie collars and successfully collected valuable movement and fate data (Brown *et al*. 1998; Chambers *et al*. 2000; Krebs *et al*. 1995; Mikesic and Drickamer 1992; Moro and Morris 2000; Pouliquen *et al*. 1990; Wilkinson and Baker 1988). However, as radio tracking technology advances our approach to animal welfare within scientific research requires ongoing adjustment (Lunney 2012). Both researchers and animal ethics committees must minimise any adverse effects on animals from radio transmitter attachment and radio tracking, which should in turn result in obtaining data more representative of normal animal behaviour (Casper 2009). This means the refinement of radio transmitter attachment methods needs to be species specific and the requirement for non-permanent attachments is a priority (Casper 2009; Rayner *et al*. 2022). Recapturing animals for transmitter removal is not always feasible; batteries can fail, and some animals may move long distances. Attaching transmitters directly to animals with adhesive glue is a non-permanent alternative, however longevity of attachment is often shorter than transmitter battery life limiting data collection (Boonstra and Craine 1986; Coetsee *et al*. 2016; O’Mara *et al*. 2014).

Electronically activated release mechanisms in radio tracking collars are available for larger mammal species, however weight and size restrictions mean these are often not suitable for smaller mammals (Buil *et al*. 2019; Cawthen and Munks 2011; Rayner *et al*. 2022). An effective non-permanent radio collar for smaller mammals relies on selecting a suitable weak-link material for incorporation into the collar, which will degrade under realistic environmental conditions, or break from sufficient force from the animal (e.g. during limb entrapment), resulting in the collar safely detaching (Cawthen and Munks 2011). Cotton and linen thread have previously been used as a weak-link in radio collars for small mammals (Cawthen and Munks 2011; Dechmann *et al*. 2014; O’Mara *et al*. 2014; O’Mara *et al*. 2019).

Weak-link radio collars have been constructed for small mammals in the 30-5000 g weight range (Cawthen and Munks 2011; Sims *et al*. 2021; Soderquist 1993) and small bats (< 45 g) (O’Mara *et al*. 2014), however no studies have focused on rodents weighing < 30 g. The small size and body weight (BW) of the house mouse (adult BW: 13-20 g) makes the selection of a suitable transmitter (<5% BW; Brooks *et al*. 2008; Buil *et al*. 2019; Coetsee *et al*. 2016; Macdonald 1978; O’Mara *et al*. 2014; Rayner *et al*. 2022) with sufficient signal range and battery life difficult, and the development of a non-permanent attachment method a further challenge. In addition to being lightweight, transmitters and their means of attachment need to be non-bulky, so they do not impede regular movement and behaviour, and robust enough to handle conspecific interactions like chewing and grooming.

Our primary aim was to identify a radio transmitter model and attachment method suitable for wild house mice living in cropping paddocks to collect movement and home range data through radio tracking studies. A radio transmitter and attachment method suitable for field deployment on mice must: (1) be of an appropriate size, shape, and weight to minimise discomfort and not compromise mouse behaviour, movement, or condition, (2) have sufficient battery life, function, and durability under field conditions for 30 days, (3) remain attached to the mouse for a minimum of 20 days and detach or show signs of detaching after 30 days, indicating that transmitters will not remain on mice indefinitely once deployed in the field, and (4) be as efficient and stress free for attachment as possible. We undertook captive trials using wild-caught house mice living in semi-natural enclosures to identify a suitable radio transmitter and attachment method for field deployment. We closely monitored how various methods affected mouse behaviour, conspecific interaction, movement, physical activity, and overall health. We further evaluated and refined our methods through radio tracking house mice in the field.

## Materials and methods

### Captive trials

From July to August 2019 and March to April 2020 we undertook captive trials at the CSIRO Crace laboratory in Canberra, Australian Capital Territory, Australia. Wild house mice live-trapped using Longworth traps (Longworth Scientific, Abingdon, United Kingdom) during separate but concurrent studies and acclimatised in the laboratory were made available for this work according to CSIRO animal ethics approvals (AEC2018-22, AEC2019-23). Mice were set up in individual ‘home’ cages (305 x 135 x 120 mm, length x width x height) containing wood-shavings, food (standard maintenance diet rat and mouse pellets, Gordon’s Specialty Stockfeeds, NSW, Australia, and commercial parrot seed mix, Watson and Williams, Australia), a water bottle, and a cardboard roll and paper towel for nesting. During the trials we tested six radio transmitter models, two attachment methods (adhesive and collar), three adhesive types for glue-on transmitters, two collar materials, and four weak-link threads for incorporation into a collar (Tables 1 and 2). We then further tested and refined the optimal adhesive and collar materials during the field sessions.

**Table 1.** Comparison of radio transmitter models

| Transmitter manufacturer | Transmitter model | Transmitter attachment type | Transmitter weight (g) | Transmitter dimensions (mm) | Transmitter antenna length (mm) | Battery life <sup>A</sup> specified by manufacturer (days) |
| --- | --- | --- | --- | --- | --- | --- |
| Advanced Telemetry Systems (ATS) | A2426 | Glue-on | 0.57 | 14 x 6 x 3 | 95 | 36 |
| Advanced Telemetry Systems (ATS) | M1410 | Collar | 0.56 | 15 x 6 x 5 | 95 | 36 |
| Holohil Systems | BD-2C | Collar | 0.94 | 11 x 6 x 4.6 | 110 | 28 |
| Holohil Systems | BD-2 | Glue-on | 0.84 | 14 x 6.5 x 3.5 | 110 | 28 |
| Lotek | PicoPip Tag Ag317 | Collar | 0.38 | 14 x 6 x 3 | 200 | 28 |
| Lotek | PicoPip Tag Ag379 | Glue-on | 0.42 | 14 x 6 x 4 | 200 | 35 |
| Lotek | PicoPip Tag Ag379 (customised<sup>B</sup>) | Collar | 0.42 | 14 x 6 x 4 | 100 | 35 |

<sup>A</sup>For a standard pulse rate range of 30-40 pulses per minute. <sup>B</sup>Transmitter-base curved and antenna length reduced from standard manufacture.

**Table 2.**
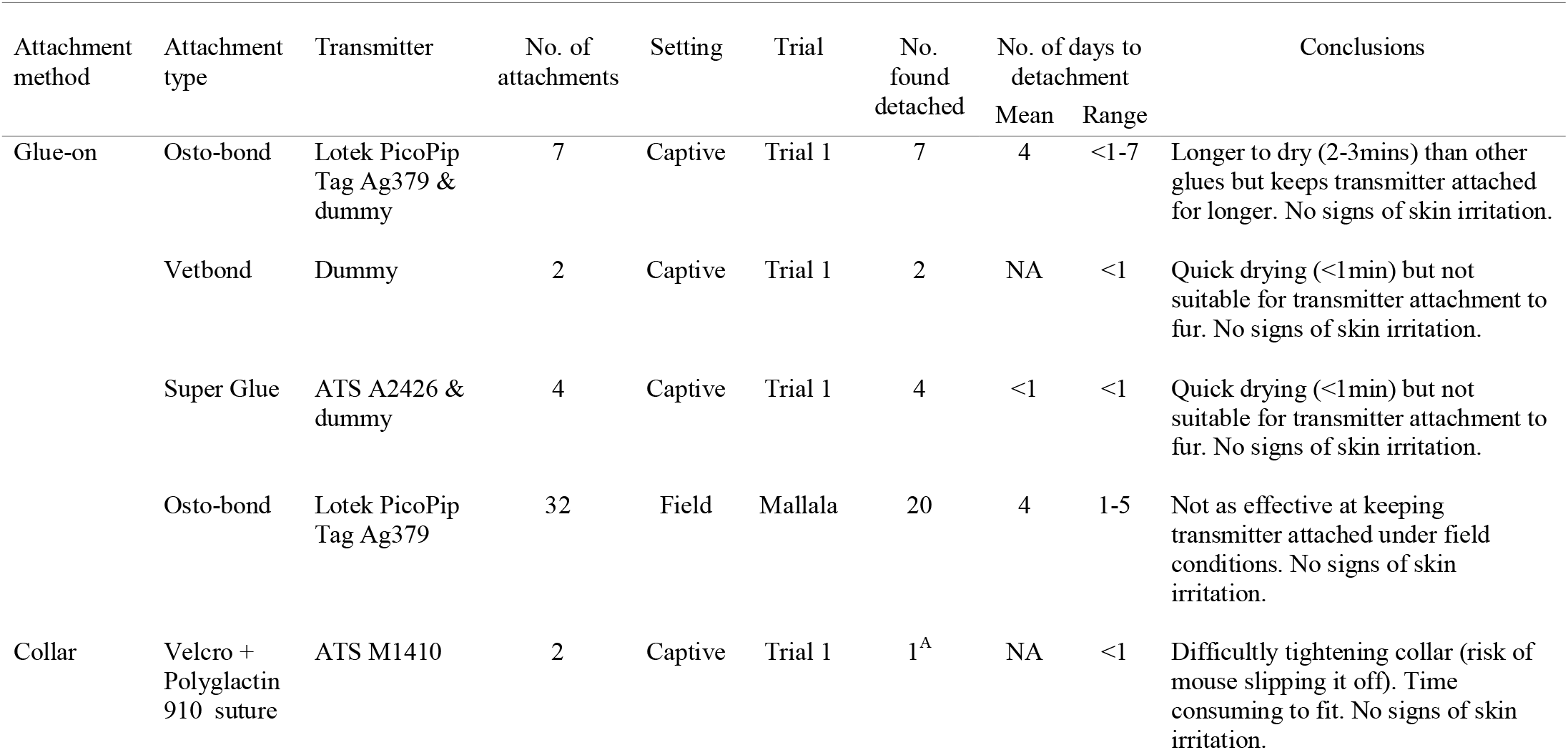
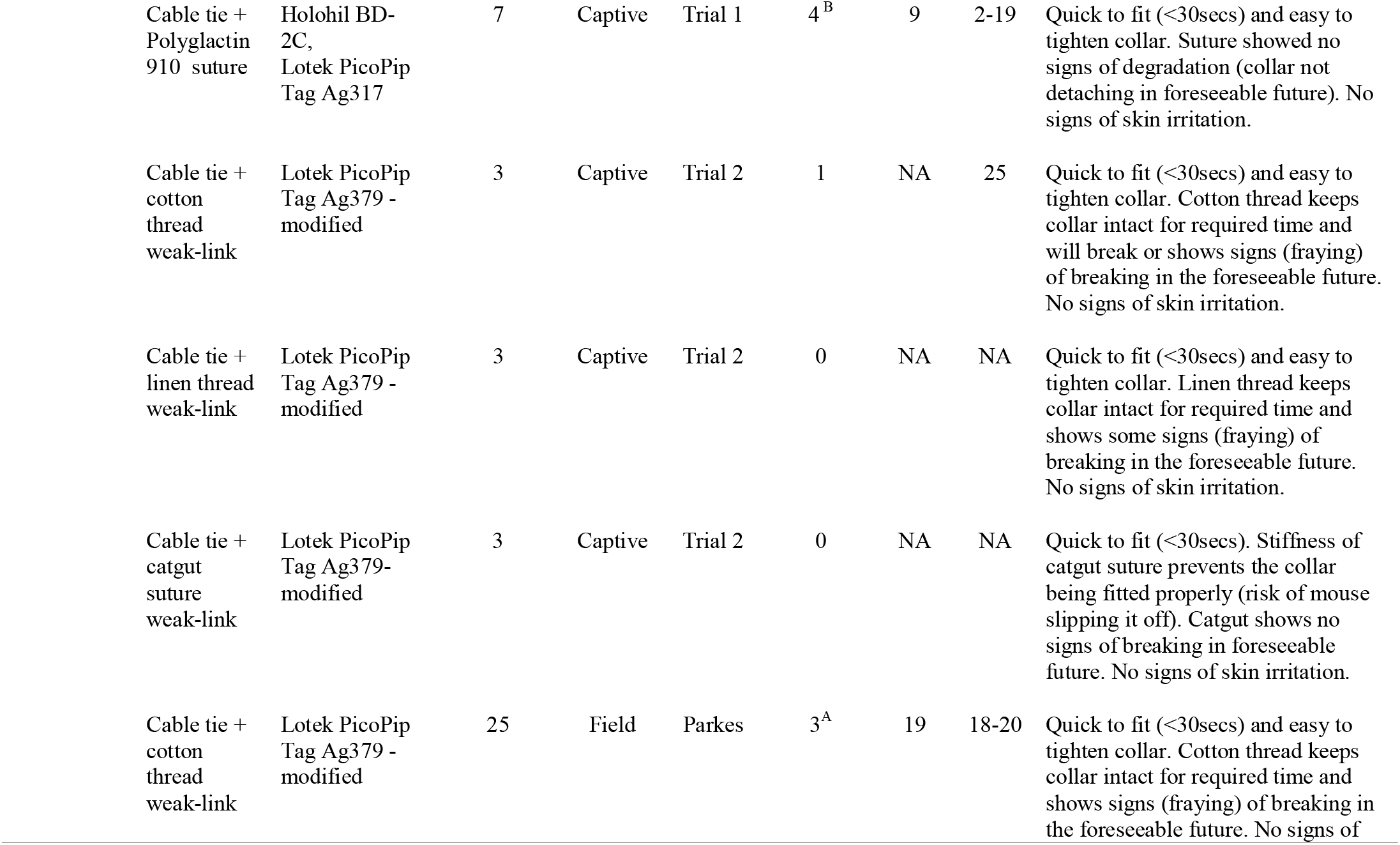
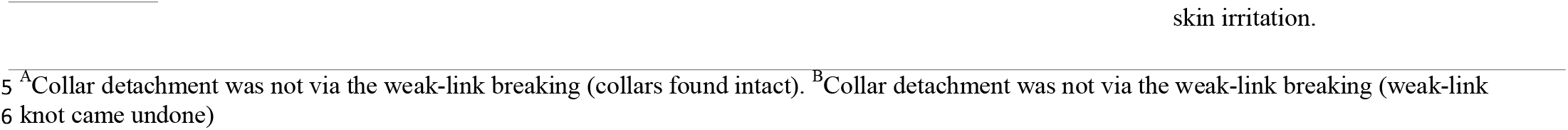
Comparison of radio transmitter attachment methods for wild house mice

For captive Trial 1, mice were placed in pairs into large enclosures with mesh lids (plastic tub: 1300 x 600 x 600 mm or glass arena: 1800 x 600 x 600 mm) to ascertain how transmitters and attachment methods would fair with conspecific interaction, particularly chewing. For Trial 2 mice were housed individually in smaller plastic Nally tubs (597 x 362 x 381 mm) with built-in mesh lids. In both trials, enclosures were set up to mimic a crop paddock with 15 cm deep soil, wheat or grass plants and straw. Wheat grain was also scattered and buried in the soil to encourage mice to search for food. Enclosures contained a water bottle, maintenance pellet food, paper towel and cardboard rolls for artificial nests. Mice were acclimatised in their enclosures for 2-3 days before radio transmitter attachment. Both trials lasted for 30 days, and mice were monitored twice daily, food and water checked daily, and weighed weekly. Mice were able to dig, run and climb in their enclosures so radio transmitters could be subjected to ‘as natural as possible’ wear-and-tear.

#### Radio transmitter models

Small and lightweight radio transmitters (<5% BW of a 13-20 g adult wild house mouse) with sufficient battery life (~30 days of transmission required) were trialled (Table 1). In Trial 1 we attached three transmitters to mice via a weak-link collar and three with adhesive (Table 2). All transmitters functioned throughout the trial, however the antennas of most were chewed partially or completely off by the end. In Trial 2 Lotek PicoPip Tag Ag379 transmitters were customised from standard manufacture for use in a collar (Table 2).

#### Attachment methods

We trialled radio transmitter models and ‘dummy’ transmitters attached to mice via weak-link collar or adhesive. Dummy transmitters were made of polymer clay encased in heat shrink and were of a similar size and weight to the real transmitters. Two collar materials and three adhesives were initially trialled with the either real or dummy transmitters on twelve mice (5M:7F, Table 2).

#### Adhesives

We trialled Osto-bond (a skin bonding latex adhesive; Montreal Ostomy, Canada), Vetbond (a tissue adhesive; 3M, Japan) and Super Glue (cyanoacrylate; Selleys, Australia) for gluing transmitters on mice (Table 2). We applied a small amount of adhesive (<10 mm diameter) to a patch of fur on the mouse’s neck just above the shoulder blades and held the base of the transmitter/dummy in place for 1-2 minutes to allow the adhesive to dry. The mouse was released into an empty holding tub for monitoring for 10 minutes to ensure the adhesive had dried and the mouse could move normally. We then replaced the mouse into its trial enclosure. The most suitable adhesive was subsequently used in field trials.

#### Weak-link collars

Nylon cable tie (2.5 x 100 mm, width x length; Crescent, Taiwan) and soft Velcro (Whites Hook & Loop Plant Tie, Australia) collars were initially trialled with an incorporated weak-link of ‘Polyglactin 910’ braided degradable suture thread (Ethicon^®^ Vicryl Rapide^™^ 4-0, USA) (Table 2). This suture was chosen for its stated rapid degradation time; designed to fall off in 10-14 days with a 0% tensile strength at 2 weeks (Byrne and Aly 2019). We then trialled three replicates of linen thread (Hemline, Australia), cotton thread (C Ne 50, Gutermann, Germany) and chromic catgut absorbable suture (USP 2/0, Bainbridge, Australia) as weak-links in cable tie collars (Table 2). Chromic catgut sutures have not been used as a weak-link in a collar however were selected for their reported loss of tensile strength after 20 days (Rengasamy and Ghosh 2010).

Cable tie collars were constructed by cutting the cable tie in half 10 mm from the clasp and smoothing the cut-ends with a file. A pin was used to make two parallel holes (1 mm apart) at both cut-ends and a needle with the weak-link material was threaded through the holes of both pieces to re-join the cable tie ends. A surgeon’s knot was tied on the top (smooth-side) of the cable tie to complete the weak-link. Transmitters were attached with 8 mm wide heat shrink tubing (Figure 1*a*). The cable tie collar weighed 0.23 g, and with transmitters attached weighed between 0.68 g (with Lotek PicoPip Tag Ag317) and 1.15 g (with Holohil BD-2C).

**Figure 1.**
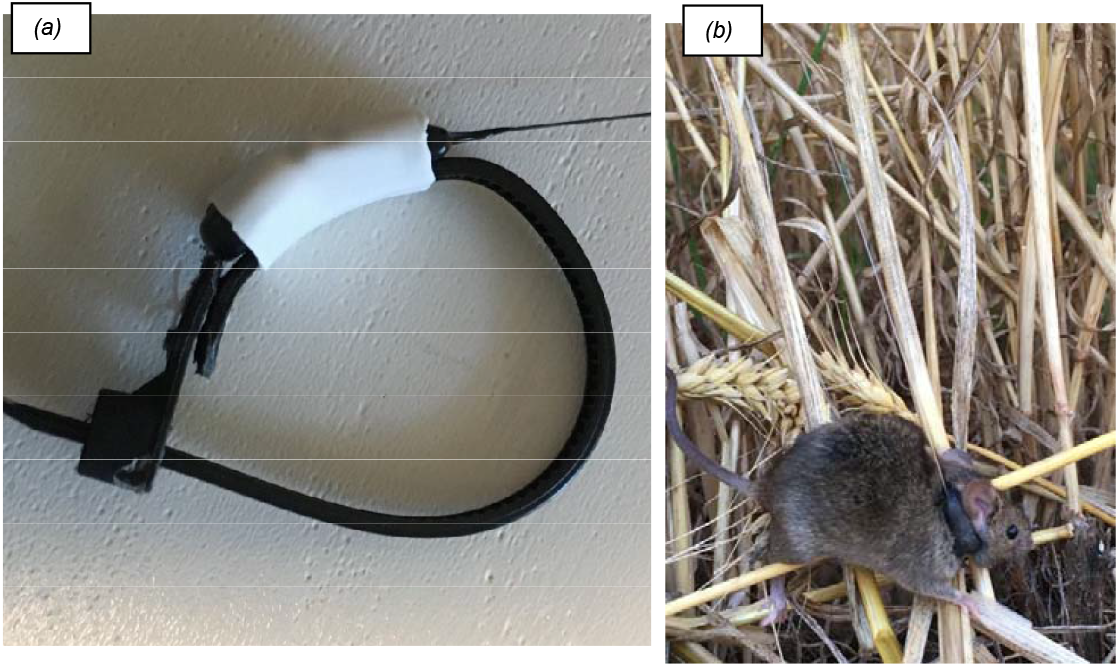
(*a*) cotton thread weak-link cable tie collar with a PicoPip Tag Ag379 attached via white heat shrink, (*b*) house mouse wearing a weak-link cable tie radio collar prior to field release. Photo credits: (a) Freya Robinson and (b) Wendy Ruscoe.

We constructed Velcro collars by cutting the Velcro into 4 mm wide x 100 mm long strips. The strip was cut in equal halves and re-joined using Polyglactin 910 suture tied off with a surgeon’s knot on the rougher side of the Velcro. Transmitters were sown onto the Velcro with cotton thread. The Velcro collar weighed 0.11 g and 0.68 g with an ATS M1410 transmitter.

We attached collars by making them into a loop, slipping them over the mouse’s head, fitting around the neck and carefully tightening. Mice were placed in an empty holding tub and monitored for 30 minutes to ensure correct collar fit and normal movement. The excess end of the collar was trimmed off and the mouse released back into its home cage. Mice were observed twice daily throughout the trial to monitor wellbeing and correct collar fit, and to check whether the collar had fallen off or the weak-link broken. Mice were weighed and the site of the collar checked weekly.

At the conclusion of the trial all weak-links were examined under a microscopic camera for signs of degradation or breakage. The preferred radio transmitter, collar and weak-link material were used in the field trials.

### Field trials

#### Field trial of adhesive

During two sessions (November and December 2019), a total of 32 house mice were fixed with glued-on radio transmitters at four sites near Mallala (34°26□18□S 138°30□35□E) in South Australia, Australia, in conjunction with another field trial where mice were being live trapped. Longworth traps containing wheat grain were set at wheat paddock sites and checked each morning at first light. Mice were weighed and given a Passive Integrated Transponder (PIT) tag (Mini HPT8, Biomark, USA) to identify individuals. Osto-bond was applied to the base of a PicoPip Tag Ag379 positioned just above the shoulder blades of the mouse and held in place for 1-2 minutes for the adhesive to dry (Table 2). The mouse was placed into a holding tub for 10 minutes to ensure the transmitter did not impede regular movement before release at its capture location. Mice fitted with transmitters weighed a mean of 18 ± 2.6 g (s.d.) and 16.3 ± 3.5 g (s.d.) for the two trapping sessions, so the transmitter weight represented means of 2.3% and 2.6% of average body weight.

**Table 3.**
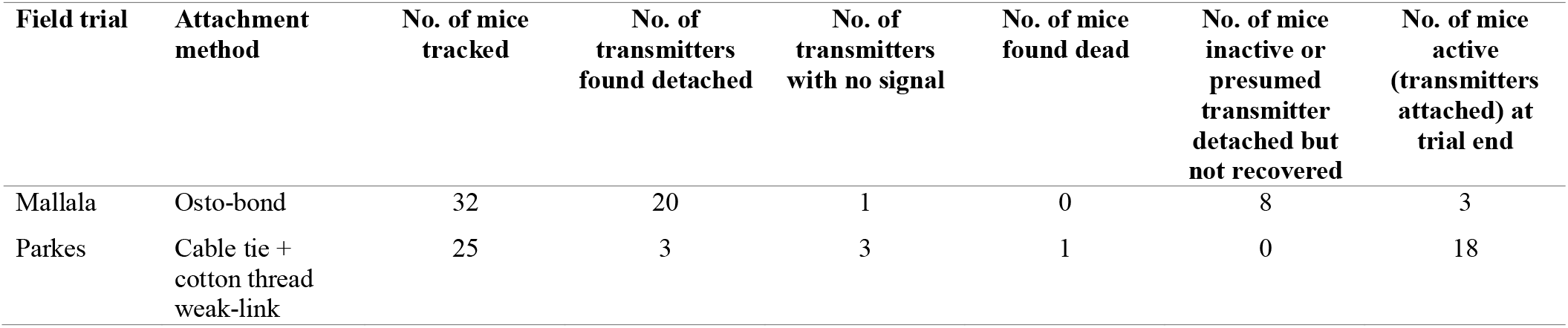
Fates of radio transmitters attached to mice during field trials

#### Field trial of weak-link collar

In November 2020 radio transmitters were fitted to wild-caught house mice at four sites near Parkes (33°8□14□S 148°10□28□E) in New South Wales, Australia, and radio tracked immediately before crop harvest and then again after harvest in December 2020. Mice were captured using Longworth traps set as described above as part of another field study. We fitted the preferred cable tie/weak-link radio transmitter collars to 25 mice (16M:9F, Table 2) weighing a mean of 18.4 ± 3.2 g (s.d.); collar and transmitter representing a mean of 3.8% of average mouse body weight. Mice were transported in traps to the nearby field house for collar attachment and monitoring. Cable tie collars with a weak-link and transmitter were constructed as per the methods described in the laboratory trial. Mice were lightly anaesthetised via inhalation of isoflurane as described by protocol (CSIRO SOP 2020-004: Anaesthesia of rodents with isoflurane in the field, AEC approved 26/5/2020) to minimise handling stress and ensure collars were fitted correctly. The collar was slipped over the mouse’s head and carefully tightened notch by notch. The process took less than 30 seconds, by which time mice had fully recovered from light anaesthesia. Mice were placed into a holding tub and monitored for 30 minutes to ensure correct collar fit and normal movement before the excess end of the collar was trimmed. Mice were then held in individual cages (305 x 135 x 120 mm) containing tissue, a toilet paper roll, apple piece and wheat grains for monitoring. Mice were released at their capture location at 2000hrs (Figure 1*b*).

#### Radio tracking

To ascertain if the radio transmitters remained attached and working, we used a Biotracker VHF Receiver and Liteflex 3-Element Yagi Antenna (Lotek, New Zealand) to locate mice. Mice were initially located from their release location and then subsequent locations between 2000hrs and 0100hrs. We estimated mice were less than three metres away when a VHF signal strength of above 90% was achieved. We confirmed this method as accurate as mice with transmitters were sighted on several occasions, at which the signal strength was over 98%. A single location fix was taken during daylight hours to establish the home burrow locations of mice. On the final day of tracking, we searched for stationary transmitters which we suspected had detached from mice.

## Results

### Transmitter type

The Lotek PicoPip Tag Ag379 was the most suitable radio transmitter. It was light (0.42 g), non-bulky and was able to be modified to better fit onto the collar. Battery life was longer than the Holohil Systems transmitters and equivalent to the ATS transmitters.

### Adhesive attachment

The adhesives we tested in the captive trial did not keep transmitters/dummys attached to mice for the minimum 20 days required. Vetbond and Super Glue kept transmitters attached for less than one day (Table 2). Osto-bond kept transmitters attached for up to seven days however attachment duration was variable (Table 2). All mice with glue-on transmitters maintained their body weight and were observed behaving and moving normally throughout the trial. Fur was pulled from the adhesive site when transmitters detached, however, we observed no skin irritation and fur grew back within a month.

We recovered 20 of the 32 Osto-bond glued-on transmitters on the ground surface or in mouse burrows within five days of attachment during the Mallala field trial (Table 3). From moving location fixes we determined only three mice retained transmitters by the end of the session (after the fifth night of tracking). We assume the remaining eight transmitters we were unable to retrieve, but still could locate signal for, had detached underground as they remained stationary for several days (Table 3).

### Weak-link materials

In the captive trial we found four of the seven Polyglactin 910 suture weak-link cable tie collars detached from mice between 2-19 days due to unwinding of the suture knot (Table 2). Collars did not detach from degradation of the suture itself. The other three collars were removed from mice; two collars were removed due to mice repeatedly slipping them off over their heads (not tight enough) and the other was removed at the end of the trial (30 days) (Table 2).

Cotton used as a weak-link was the only material that broke resulting in the cable tie collar detaching from a mouse after 25 days (Table 2). On several occasions we found that some collars with intact catgut weak-links had fallen off mice. Catgut was not flexible like the linen and cotton threads which meant the cable tie collar was more rigid at the weak-link point and did not fit as snuggly around the mouse’s neck. This resulted in mice being able to slip the catgut weak-link collars over their heads. The collars with linen weak-links remained on mice for the 30 days of the trial.

On examination of weak-link material under a microscopic camera after 30 days, both cotton and linen threads showed signs of deterioration, whereas catgut did not (Figure 2). The two intact cotton weak-links were frayed and most strands were severed from rubbing against the cable tie, suggesting the collars probably would not have stayed on the mice much longer than the 30 days. The linen weak-link was less degraded, with fewer strands of the thread severed and frayed. The catgut weak-links showed no signs of degradation and the catgut was intact, suggesting collars would probably not have detached in the foreseeable future.

**Figure 2.**
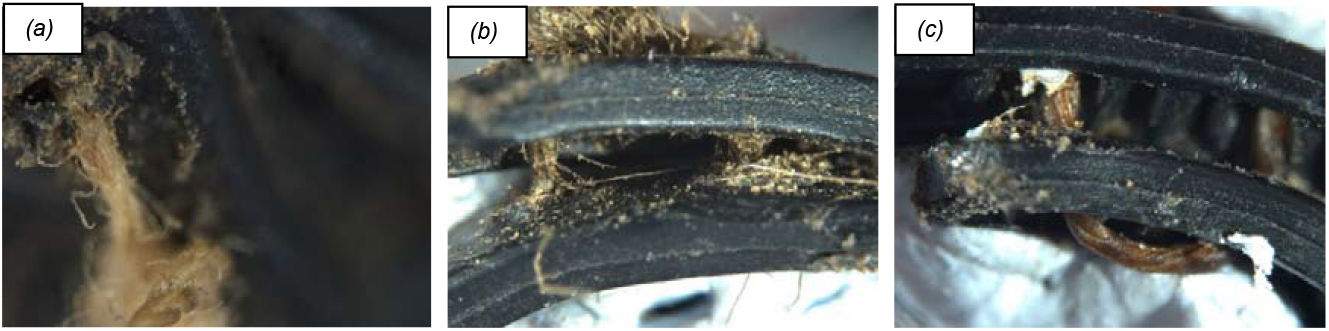
Images of weak-link materials in cable tie collars photographed under a microscope camera after removal from mice at day 31 of captive laboratory Trial 2: (*a*) fraying cotton thread weak-link showing only a few individual strands holding the collar together, (*b*) fraying linen thread weak-link showing a few severed strands only, (*c*) catgut suture weak-link showing no degradation. Photo credit: Freya Robinson.

### Collars

Attachment of the Velcro collar was difficult as tightening the collar to fit on the mouse was very awkward and time consuming (taking up to five minutes). For this reason, only two Velcro collars were attached to mice and neither remained on for more than one day as the collar could not be tightened enough to prevent it slipping it off (Table 2). The Velcro proved unsuitable in the laboratory, so was not trialled further.

We found the cable tie collars quick to fit in the laboratory, though ensuring the collar was sufficiently tightened was at times challenging due to mice squirming or hunching during fitting. It was essential to ensure collar tightness as on several occasions’ mice slipped their forelimbs under the collar trying to remove it. By monitoring mice for 30 minutes after collar fitting, we avoided mice injuring themselves or getting their forelimbs permanently stuck. We observed a slight decrease in activity (running and jumping) in some mice immediately after collar fitting, however all resumed normal activity once returned to their home trial cages. We observed mice exploring the surroundings in their trial tubs, running, climbing, jumping, digging, grooming, feeding and burrow-digging. Mice maintained condition and body weight throughout the trials. Mice had minor fur loss, and a few had minimal skin irritation around the neck after collar detachment and/or removal.

During the Parkes field trial we retrieved three collars which had come off mice, however none had detached because the cotton thread weak-link had broken; two collars were found intact (likely slipped over the mouse’s head) at 19 and 20 days after attachment, and the other collar’s weak-link knot had come undone and was found 18 days after attachment (Table 2). On the fifth and final day of radio tracking (18-21 days after collar attachment) we concluded that 18 of the 25 mice (72%) were active and still wearing functioning transmitters (Table 3). We found one dead mouse which had been flattened on a wheel track most likely during harvest and could not pick up the signals of three transmitters (Table 3).

Over the course of the trial we sighted five mice with transmitters and recaptured three mice in our live capture traps. We examined the conditions of the recaptured mice and only some minor fur loss around the neck from the collar was evident. Most of the transmitters we retrieved or sighted on recaptured mice had aerials chewed fully off however the transmitter itself remained intact and well attached to the collar. Despite this we could pick up the signal of most transmitters from distances of up to 50 m. The cotton weak-links were also showing signs of fraying. A high proportion of collars remained on mice at the end of the study (18-21 days after collar attachment) which allowed sufficient time for meaningful home range locations to be collected.

## Discussion

Trialling radio transmitters and attachment methods on captive house mice under semi-natural laboratory settings allowed us to identify not only suitable methods prior to field deployment but also identify and solve challenges before they became problems in the field. The adhesives we trialled were not suitable for attachment of radio transmitters to mice under field conditions. Vetbond and Super Glue were abandoned early in the laboratory trial when transmitters detached from mice within a day. Transmitter attachment duration with Osto-bond was highly variable in the laboratory (<1-7 days), however we found this was further reduced under field conditions (1-5 days). Wilkinson and Baker (1988) also had variable success using colostomy adhesive with transmitters detaching from house mice between two to 17 days. Krebs *et al*. (1995) found attaching transmitters to house mice with skin glue was unsuccessful. We suggest Osto-bond is the best option for studies intending to track juvenile or small mammals over several days, where cable tie collars are unsuited.

From our laboratory trials we identified a customised Lotek PicoPip Tag Ag379 as the most suitable radio transmitter for a house mouse and our required minimum 20 days of data collection. This transmitter is light and non-bulky and can be glued-on or incorporated into a collar. Cable tie collars with a built-in cotton thread weak-link were the preferred method of attachment under field conditions. They remained on mice long enough to collect radio tracking data between field sessions. Radio collars retrieved between 18-20 days post deployment showed signs of the cotton thread fraying indicating collars likely would not have remained on mice indefinitely in the field if the animals could not be re-trapped, satisfying animal welfare concerns.

We are not aware of any studies which have trialled or used a weak-link in radio collars for small rodents (<30 g). In a review of radio collar attachment methods for Australian mammals weighing between 35-5500 g, Rayner *et al*. (2022) found the effectiveness of weak-links was highly variable as similar materials degraded at different rates from varying environmental conditions. For this reason Coetsee *et al*. (2016), Sims *et al*. (2021) and Rayner *et al*. (2022) all recommend designing weak-link collars specifically for the target species and trialling in captive conditions resembling field conditions until a safe and effective weak-link collar is identified. Our laboratory trials quickly identified catgut and Polyglactin 910 suture as unsuitable weak-links for house mice living in cropping paddocks. Cotton thread was deemed most suitable in our laboratory trial when a collar detached at 25 days. We anticipate that collars would not have remained on mice in the field indefinitely as fraying of the cotton thread was evident in collars that were retrieved.

Non-release nylon cable ties collars have been used in radio tracking studies of house mice (Chambers *et al*. 2000; Krebs *et al*. 1995; Moro and Morris 2000) and other small rodents weighing less than 30 g (De Mendonça 1999; Gray *et al*. 1998; Tew and Macdonald 1993). We found the collars had no detrimental effects on house mouse condition or behaviour during our laboratory trials or in those mice we resighted or recaptured in the field. Cable tie collars are light, quick to fit, easily tightened and were able to have a weak-link readily incorporated.

Few studies have evaluated the effects of radio collars on house mice and other small rodents (Ormiston 1985; Pouliquen *et al*. 1990; Wolton and Trowbridge 1985). This likely reflects the minimal monitoring of animals post transmitter attachment and low recapture rates in the field, as well as a lack of captive trials prior to field deployment. Pouliquen *et al*. (1990) monitored the effects of radio collars on wild house mice in the laboratory showing decreased physical and social activity immediately after collar fitting but a return to normal activity after four hours. In both our laboratory trials we also observed a temporary decrease in normal activity (running and jumping) in some individuals within the 30 minutes of monitoring post collar attachment, however all mice resumed normal activity after being placed back into their home tubs. Radio collars were not reported as adversely affecting feeding behaviour and social interactions in wood mice (*Apodemus sylvaticus*) (Wolton and Trowbridge 1985) and meadow voles (*Microtus pennsylvanicus*) (Webster and Brooks 1980), and no adverse effects on reproductive effort, home range size and movement was reported for collared white footed mice (*Peromyscus leucopus*) in the field (Ormiston 1985). Few studies have reported injury or loss in body condition due to radio collars (Ormiston 1985; Pouliquen *et al*. 1990). De Mendonca (1999) also determined that the shape and size of the radio collar package rather than the weight had a greater impact on yellow-necked mice (*Apodemus flavicollis*).

Choosing a suitable attachment method and duration for a species is just as important from an animal welfare perspective as focusing on percentage body weight (Lunney 2012). Casper (2009) suggested that the 5% or 3% body weight rule for transmitter attachment was too simplistic and attachment methods need to be designed specifically for each species and situation, with some aspect of monitoring or pilot studies prior to field deployment. Our laboratory trial led us to request modified Lotek PicoPip Tag 379 transmitters that were slightly moulded to the shape of the collar.

We found monitoring mice with collars for several hours before their release back into the field was very important for mouse welfare. Some mice would attempt to remove the collar, which on a few occasions resulted in entrapped forelimbs. We were able to promptly rectify this and tighten the collar to avoid further limb entrapment, which would have caused death in the field. Most radio tracking studies of house mice or small rodents do not report animal monitoring after radio collar fitting. For those that do, collared animals were held and observed for 3-10 minutes or ‘several’ minutes (De Mendonça 1999; Mikesic and Drickamer 1992; Tew and Macdonald 1993), for no more than one hour (Banks *et al*. 1975; Ormiston 1985) and for up to 2-6 hours in captivity (Gray *et al*. 1998; Moro and Morris 2000). Although no optimal monitoring time is discernible from these studies, shorter periods rely on the collar being fitted accurately once. We found this difficult to consistently achieve, especially on squirmier mice and therefore recommend a longer observation time. The use of an anaesthetic may increase the accuracy of fitting collars, reduce animal stress and potential injury during fitting and minimise the time taken to accurately fit the collar. De Mendonça (1999) suggested that the stress associated with being handled and fitted with a collar may be more important to animal wellbeing than any effects from wearing the collar. Numerous studies have used light anaesthesia to aid in fitting collars and reducing stress to small rodents (Hamley and Falls 1975; Mineau and Madison 1977; Tew and Macdonald 1993). We found fitting collars to lightly anaesthetised mice was preferable as we were able to properly fit collars without prolonged handling.

The collection of meaningful radio tracking data relies on careful selection of a radio transmitter and attachment method suitable for the study species. By trialling various radio transmitter models and attachment methods prior to field deployment we identified a suitable cotton thread weak-link cable tie collar and transmitter for radio tracking house mice in cropping paddocks. The development of this weak-link radio collar is an important improvement on existing permanent radio collars for small mammals weighing < 30 g.

## Acknowledgements

We sincerely thank the farmers for allowing us to undertake this experiment on their properties. We thank Leigh Nelson and Ken Young (GRDC) for ongoing support. FR and WAR wrote the first draft manuscript. All authors collected experimental data and contributed to the final manuscript.

## Ethics Approval

This study was approved by the CSIRO Wildlife and Large Animal, Animal Ethics Committee (Approval numbers: 2019-18, 2019-24, 2019-31) and adheres to the 8th Edition of the Australian Code and Use of Animals for Scientific Purposes. This article does not contain any studies with human participants performed by any of the authors.

## Availability of data and material

Data are held by the Commonwealth Science and Industry Research Organisation (CSIRO), Australia, and may be made available upon reasonable request to the senior author.

## Supplementary information

None

